# Mitochondrial Transplantation in the Eye: A Review and Evaluation of Surgical Approaches

**DOI:** 10.64898/2026.04.06.716722

**Authors:** Bertan Cakir, Tsai-Chu Yeh, Cheng-Hui Lin, Man-Ru Wu, Éric Boilard, Martin Pelletier, Aneal M Singh, Yann Breton, Samir Patel, Tom Benson, David RP Almeida, Sui Wang, Vinit B. Mahajan

## Abstract

**Purpose:** Mitochondrial dysfunction contributes to major blinding diseases, including age-related macular degeneration and glaucoma. Although mitochondrial transplantation has shown therapeutic potential in multiple organ systems, translation to the eye remains limited, partly due to uncertainty regarding optimal delivery. We summarize the biologic rationale and preclinical evidence supporting ocular mitochondrial transplantation and present feasibility data evaluating clinically relevant delivery routes.

**Methods:** We conducted a focused narrative review of ocular mitochondrial transplantation. For feasibility experiments, mitochondria with an endogenous fluorescent dye were isolated from liver donor mice. Postnatal day 7 pups received subretinal injections, and adult CD1 mice received intravitreal injections, including optic nerve head directed delivery. Eyes were analyzed using fluorescence microscopy and immunohistochemistry. Mitochondrial uptake was assessed in cultured retinal pigmental epithelial (RPE) cells using co-incubation assays. Suprachoroidal delivery feasibility was evaluated in cadaveric human near-real surgical specimens using a novel dedicated suprachoroidal injector.

**Results:** The literature on ocular mitochondrial transplantation remains limited and consists primarily of small preclinical studies using intravitreal delivery and imaging-based detection. In our experiments, intravitreal delivery produced donor signals predominantly within inner retinal layers, with enrichment along retinal nerve fiber bundles when directed toward the optic nerve head. Cultured RPE cells demonstrated dose-dependent uptake of exogenous mitochondria. Subretinal delivery localized donors signal to the RPE and adjacent outer retina. Suprachoroidal injections demonstrated procedural feasibility with reliable access to the suprachoroidal space and visible injectate distribution.

**Conclusions:** Ocular mitochondrial transplantation is in an early stage of investigation. Our feasibility data indicate that established posterior-segment delivery routes expose distinct retinal compartments and that route selection strongly influences anatomic distribution. Further studies are needed to verify intracellular uptake, define dosing and durability, and evaluate safety in disease-relevant models.

## Introduction

Mitochondria are essential for maintaining cellular energy and human health. They play critical roles in fundamental cellular processes, including energy production via oxidative phosphorylation, amino acid and lipid metabolism, heme biosynthesis, the modulation of immune responses, and the regulation of apoptosis.^1–3^ Dysfunction of these organelles is implicated in a wide spectrum of human diseases. The retina and retinal pigment epithelium (RPE) are among the most metabolically active tissues in the human body.^4^ These tissues rely heavily on mitochondrial function for phototransduction, phagocytosis of photoreceptor outer segments, and maintenance of ion gradients required for vision. Consequently, mitochondrial dysfunction disproportionately affects the visual system and contributes to a range of ocular pathologies.

Mitochondrial function declines with age, and mitochondrial dysfunction has been identified as one of the hallmarks of aging.^5^ This decline has been linked to various age-related diseases, including Alzheimer’s disease, Parkinson’s disease, age-related macular degeneration (AMD), and glaucoma.^6,7^ Additionally, mitochondrial dysfunction plays a significant role in other vision-threatening conditions, such as inherited mitochondrial diseases, including Leber hereditary optic neuropathy and dominant optic atrophy, as well as metabolic disorders like diabetic retinopathy.^8–10^

Given the central role of mitochondrial dysfunction in major blinding conditions and aging, improving mitochondrial function represents a compelling therapeutic strategy.^11^ Consequently, there has been growing interest in developing novel therapeutic strategies aimed at improving mitochondrial function.^12–18^ While significant progress has been made, the complexity of mitochondrial biology poses considerable challenges for drug-based therapies, leading to limited treatment responses.^19^ Transplantation of intact, healthy mitochondria has therefore emerged as a promising alternative strategy that may augment cellular bioenergetics more directly.

Despite growing momentum for mitochondrial transplantation in systemic disease models, its application to ophthalmology remains at an early stage, in part because effective delivery to specific ocular compartments is a fundamental hurdle. To advance mitochondria-based therapies for eye diseases, it is essential to determine whether mitochondria can be delivered to key targets including the neural retina, optic nerve head, and RPE. Here, we review the therapeutic landscape of mitochondrial transplantation and present new feasibility data assessing delivery of isolated mitochondria to the retina, optic nerve head, and RPE via subretinal and intravitreal injection in mice, and via a novel suprachoroidal delivery system in ex vivo human surgical specimens.

### The Therapeutic Landscape of Mitochondrial Transplantation

#### Intercellular Mitochondrial Transfer and Cellular Uptake

Mitochondria are thought to have originated from a distant endosymbiotic event in which a proto-eukaryotic cell fused with an α-proteobacterium, enabling the high metabolic capability that characterizes modern eukaryotic life.^20^ For many years, mitochondria were thought to be inherited exclusively during cell division (vertical transmission). However, research over the past two decades has demonstrated that mitochondria can also move horizontally between neighboring cells, a process now broadly termed intercellular mitochondrial transfer. The first functional evidence showing this process came from a co-culture model in which mtDNA-deficient ρ cells regained aerobic respiration after acquiring mitochondria from adjacent cells.^21^

Several mechanisms of horizontal transfer have since been identified, including direct cell-cell connections including tunneling nanotubes (TNTs) and gap junction channels (GJCs), extracellular vesicle-associated mitochondria (EVMs), and the release of free/naked mitochondria. Among these, TNT-mediated transfer is the best characterized. TNTs are transient, cytoplasmic F-actin based bridges that physically link neighboring cells and allow the movement of mitochondria via motor proteins such as MIRO1.^22^ Mitochondrial transfer through these structures has been described in several biological systems, including cardiovascular tissues,^23–25^ the respiratory epithelium,^26,27^ immune system,^28,29^ corneal epithelium,^30^ tumors,^31,32^ and the central nervous system,^33–35^ suggesting that direct cell to cell mediated transfer is a conserved biological mechanism in various and unique circumstances. Another major route of mitochondrial transfer occurs through EVMs, where cells package mitochondrial material into vesicles that range from small tetraspanin-positive particles that primarily carry fragmented damaged mitochondrial components ^36,37^ to larger LC3-positive vesicles that contain intact mitochondria derived from the autophagolysosomal system.^38^ Although this pathway is most often described as a mechanism for clearing damaged mitochondria during cellular stress, recent studies have shown that EVMs can also mediate transfer of intact, functional mitochondria capable of improving bioenergetic function in recipient cells. For example, thrombin-activated platelets can deliver respiration-competent mitochondria to neighboring cells ^39^, and astrocyte-derived mitochondria can support hypoxic neurons after stroke, improving their survival.^40^ Although the entire process of vesicle docking, internalization, and mitochondrial integration is not fully resolved yet, EVM-mediated transfer is increasingly recognized as a physiologically significant process.

A third mode of mitochondria transfer involves the release of free or naked mitochondria into the blood stream. Both platelets and adipocytes have been shown to contribute to the circulating mitochondrial pool, and these free mitochondria can be distinguished from vesicle-bound cargo by expression of translocase of the outer mitochondrial membrane-22 (TOM22) and lack of tetraspanins.^36,41^ Naked mitochondria (measure approximately 0.5–1 µm in size) released from thrombin activated platelets^39^ have been shown to exhibit membrane potential and retain full-length mtDNA, suggesting functional capacity.^42,43^ Their release is dependent on key mitochondrial fission regulators.^44^ Uptake into recipient cells is thought to occur through a heparan-sulfate-dependent phagocytic mechanism requiring 6-O sulfation,^45^ and one study has shown that transferred mitochondria can escape endosomes and contribute to oxidative metabolism.^46^

Taken together, these observations suggest that intercellular mitochondrial transfer is a physiologic and conserved process, enabling cells to support each other during metabolic stress or injury. This intrinsic capacity for horizontal mitochondrial transfer provides the biological foundation for the emerging therapeutic strategy of mitochondrial transplantation.

#### Functional Roles of Intercellular Mitochondrial Transfer

While we described the main routes by which mitochondria can transfer between cells, the next question is what functional impact have transferred mitochondria have in recipient cells. In the literature, the most frequent reported functional effect of mitochondrial transfer is improved bioenergetic function, which is linked to a variety of context-specific effects such as neuroprotection, wound healing, hematopoiesis, reduced inflammation, metabolic conditioning, and bone remodeling.^11^

The first described beneficial effect of mitochondrial transfer came from experiments showing that mtDNA-deficient ρL cells, which are unable to perform oxidative phosphorylation, regained aerobic respiration after acquiring mitochondria from co-cultured donor cells.^21^ Interestingly, the degree of functional benefit of transferred mitochondria appears to be context-dependent, with the biggest metabolic effect found under cellular energetic stress across multiple studies. For example, in vivo metabolically compromised peritoneal macrophages lacking the complex I subunit NDUFS4 retained injected wild-type mitochondria longer and showed a larger metabolic response than wild-type macrophages, despite similar uptake.^47^ Consistent with this stress-dependent effect, BV2 microglial cells do not substantially increase aerobic respiration after exposure to purified mitochondria under baseline culture conditions but can use exogenous mitochondria to recover respiration following pharmacologically induced mitochondrial dysfunction. ^47^ This crisis-dependent pattern is also reflected in ischemic settings where neurons and cardiomyocytes can receive mitochondria with associated improvements in local energetics and cell rescue.^40,48–50^

#### Therapeutic Applications Across Disease Models

After in vivo transplantation of isolated autologous mitochondria was first reported in 2009 by McCully and colleagues,^26^ numerous preclinical studies across multiple organ systems have been persued, including traumatic brain injury,^27^ cerebral ischemic injury,^28^ Alzheimer’s disease,^29^ Parkinson’s disease,^30^ heart failure,^31^ myocardial infarction,^32^ sepsis,^33^ acute kidney injury,^34^ and ischemia-reperfusion lung injury.^35^ Across these models, mitochondrial transfer has been associated with improved metabolic function and tissue recovery, encouraging the translation of this approach into clinical practice.

Recent research has also demonstrated the potential for mitochondrial transfer to treat genetic mitochondrial diseases. Nakai et al. have shown that both mouse- and human-derived mitochondria improved neurological function and survival in a mouse model of Leigh syndrome, supporting mitochondrial transfer as a possible therapeutic approach in primary mitochondrial conditions.^36^ With growing preclinical evidence, early clinical studies have also shown promising results including in myocardial ischemia-reperfusion injury and cerebral ischemia.

Collectively, these studies establish two key points relevant to ophthalmology. First, mitochondrial transplantation is increasingly supported as a biologic therapeutic strategy across diverse tissue types and disease mechanisms. Second, delivery modality is highly context dependent: systemic organs often permit direct parenchymal injection or vascular administration, whereas ocular tissues require compartment-specific delivery approaches that achieve local exposure without compromising delicate retinal structures.

While the benefit of mitochondrial transfer becomes increasingly clear, the mechanisms through which these benefits occur remain poorly understood. Recent work in a neurodegeneration model provides some evidence that transplanted mitochondria may functionally integrate into host neuronal networks colocalizing with endogenous mitochondria, improving cellular energetics with restored ATP levels. This effect lasted for 4 weeks coinciding with phenotypic rescue.^51^ Another mechanism contrary to the functional integration has been described in an endothelial model where exogenous mitochondria are internalized by macropinocytosis and rapidly targeted for PINK1-Parkin dependent mitophagy, triggering mitochondrial biogenesis and metabolic reprogramming even when donor mitochondria are energetically compromised.^52^

Despite these advances, the precise mechanisms mediating these functional benefits are still incompletely defined and controversial. However, it is increasingly likely that mitochondrial transfer can act through multiple pathways depending on tissue type and state, such as direct bioenergetic incorporation, mitophagy-induced signaling, and possible unknown extracellular or paracrine-like pathways.

#### Mitochondrial Dysfunction in the Eye

Mitochondrial dysfunction has emerged as a central pathogenic mechanism across major posterior-segment diseases, reflecting the exceptionally high energetic demands of the RPE, photoreceptors, and retinal ganglion cells (RGCs). These highly metabolically active cells depend on efficient oxidative phosphorylation, balanced redox homeostasis, and continuous mitochondrial quality control. When these systems fail, bioenergetic decline, oxidative injury, and metabolic uncoupling result in retinal and optic nerve degeneration. Given these vulnerabilities, mitochondrial transplantation represents a promising therapeutic strategy.

##### Age-Related Macular Degeneration

Replacing dysfunctional mitochondria in the RPE and retina may hold significant therapeutic value for age-related macular degeneration (AMD). Mitochondrial dysfunction in the RPE has been identified as one of the key drivers of AMD disease pathogenesis.^53^

Studies of human donor eyes reveal that, in AMD patients, RPE mitochondria show disrupted architecture, reduced mitochondrial mass, increased mitochondrial fragmentation, and altered expression of mitochondrial proteins, including proteins of the electron transport chain.^7–9^ Mitochondrial DNA (mtDNA) damage is thought to be one of the key upstream drivers of mitochondrial structural and functional decline in AMD and is particularly evident in the RPE, consistent with its heavy reliance on oxidative phosphorylation and associated oxidative stress. In support of this hypothesis, a study of human donor RPE from AMD patients and age-matched controls showed that mtDNA lesions preferentially accumulate with AMD progression. In the same study, mtDNA damage exceeded that of two nuclear genes with total mitochondrial genome injury estimated to be about 8-fold higher, supporting selective mtDNA vulnerability in AMD-associated RPE degeneration.^54^ Experimental evidence that mitochondrial oxidative stress can drive RPE dysfunction comes from an RPE-specific Sod2 knockout mouse model, in which loss of mitochondrial superoxide dismutase increases mitochondrial ROS. Elevated oxidative stress in vivo was sufficient to cause RPE functional impairment, accompanied by mitochondrial structural abnormalities leading to dysfunction with reduced ATP levels.^55^ This impaired energetics lead to an increased expression of key glycolytic enzymes in the RPE, suggesting a shift toward glycolysis when mitochondrial function is compromised.^55^

This glycolytic switch has important downstream consequences for photoreceptor metabolism, linking primary RPE dysfunction to broader retinal vulnerability. In the “metabolic ecosystem” model, the RPE largely spares glucose for photoreceptors, which rely on glycolysis and export lactate that the RPE in return uses for oxidative phosphorylation and ATP generation.^56^ When mitochondrial function in the RPE is compromised and the RPE shifts toward glycolysis, this coupling can break down, reducing glucose delivery to photoreceptors which leads to photoreceptor starvation and ultimately degeneration.

OCT-based analyses in humans provide complementary clinical support for mitochondrial decline in AMD. The ellipsoid zone (EZ) reflects the photoreceptor inner segments and is thought to be caused by its high mitochondria density. A longitudinal study showed that relative EZ reflectivity declines with age and more markedly with AMD progression.^57^ Although EZ reflectivity is an indirect structural metric, these data are consistent with progressive decline of photoreceptor inner-segment mitochondrial integrity in AMD, supporting the broader model that mitochondrial dysfunction is a core feature of disease pathogenesis.

Together, these observations provide a rationale for mitochondrial transplantation as a strategy to restore healthier mitochondrial populations in the RPE, improve bioenergetic function, and preserve photoreceptor health in AMD.

##### Diabetic Retinopathy

Similarly, mitochondrial dysfunction is increasingly recognized as a central driver of diabetic retinopathy (DR). Historically, DR was viewed primarily as a microvascular disease in which pericyte loss and endothelial injury destabilize capillaries, leading to microaneurysms, vascular leakage, vasoregression, and at later stages pathologic neovascularization.

A prevailing mechanistic framework underpinning this microvascular model is that diabetes amplifies oxidative stress, which results in the injury of the retinal endothelium, with mitochondria serving as a key source of reactive oxygen species.^58^. However, accumulating evidence now supports the concept that early retinal neurodegeneration can precede overt microvascular pathology. ^59–62^ These observations align with subjective decline in contrast sensitivity, color vision, and overall all quality of vision in patients with minimal to no signs of diabetic retinopathy.^63–65^ Consistent with a neuroretinal origin of oxidative stress, Du et al. reported that diabetes-induced oxidative stress arises predominantly from photoreceptors rather than endothelial cells. ^66^ This heightened oxidative burden was accompanied by inflammatory activation, which is widely regarded as a core contributor to DR pathogenesis, resulting in vascular injury and regression in the diabetic retina.^66^ This neural retina contribution is further supported by clinical and experimental observations that photoreceptor degeneration (e.g., retinitis pigmentosa) is associated with relative protection from DR severity.^67,68^

Several mechanisms may explain why diabetic photoreceptors generate excess superoxide. One plausible model is that diabetes creates a state of substrate overload with increased glucose availability and a shift toward fatty-acid oxidation driving high electron flux through mitochondrial oxidative metabolism. This increased electron pressure promotes electron leak to oxygen and superoxide generation, particularly at complexes I and III.^69^ While mtDNA appears relatively preserved in early stages of diabetes,^70^ later stages show altered mtDNA ^71^ resulting in a decline in mitochondrial transcription and secondary ETC and a cycle of increased electron leak and oxidative stress. Diabetes-induced mitochondrial dysfunction ultimately reduces ATP production, disrupts redox balance, promotes inflammation, and accelerates neuronal apoptosis, contributing to neurovascular uncoupling and ultimately capillary dropout and pathologic neovascularization.^72^ Importantly, mitochondrial oxidative stress and dysfunction persists even after normalization of blood glucose,^73^ likely due to accumulated mtDNA damage and durable epigenetic alterations. ^74^

Together, these findings highlight mitochondrial dysfunction as a critical therapeutic target to halt or slow DR progression. In this context, intravitreal mitochondrial transplantation could mitigate oxidative injury and restore metabolic balance in the neuroretina, potentially preventing early neuronal dysfunction and slowing progression of diabetic retinopathy.

##### Glaucoma and other Optic Neuropathies

In glaucoma, mitochondrial dysfunction is increasingly recognized as a central mechanism of retinal ganglion cell (RGC) degeneration.^75,76^ RGCs and their axons at the optic nerve head are highly vulnerable due to their dense mitochondrial populations and reliance on oxidative phosphorylation.^77^ Experimental animal studies demonstrate early impairment of mitochondrial axonal transport following ocular hypertension which is exacerbated by aging. Brief elevations in intraocular pressure in primates lead to decreased axonal mitochondrial transport resulting in swollen dysfunctional mitochondria at the lamina cribrosa.^78,79^ Furthermore, inherited mitochondrial optic neuropathies, such as Leber hereditary optic neuropathy and dominant optic atrophy which selectively affect RGCs, further highlights the vulnerability of RGC nerve fibers with mitochondrial dysfunction.^6^ Impaired axonal transport, vascular dysregulation, and decreased mitochondrial capacity at the optic nerve head collectively render this region susceptible to secondary insults such as aging, nocturnal hypotension, and obstructive sleep apnea. Targeting mitochondrial dysfunction has great potential to slow glaucoma progression and prevent vision impairment. By replenishing damaged RGC mitochondria and restoring axonal energy capacity, mitochondrial transplantation holds promise for protecting RGCs and preserving vision in glaucoma.

#### Mitochondrial Transplantation in the Eye

Ocular data on mitochondrial transplantation remain limited, and much of the current evidence comes from in vitro systems and small preclinical in vivo studies (**Figure 1, Table 1 and 2)**

**Figure 1.**
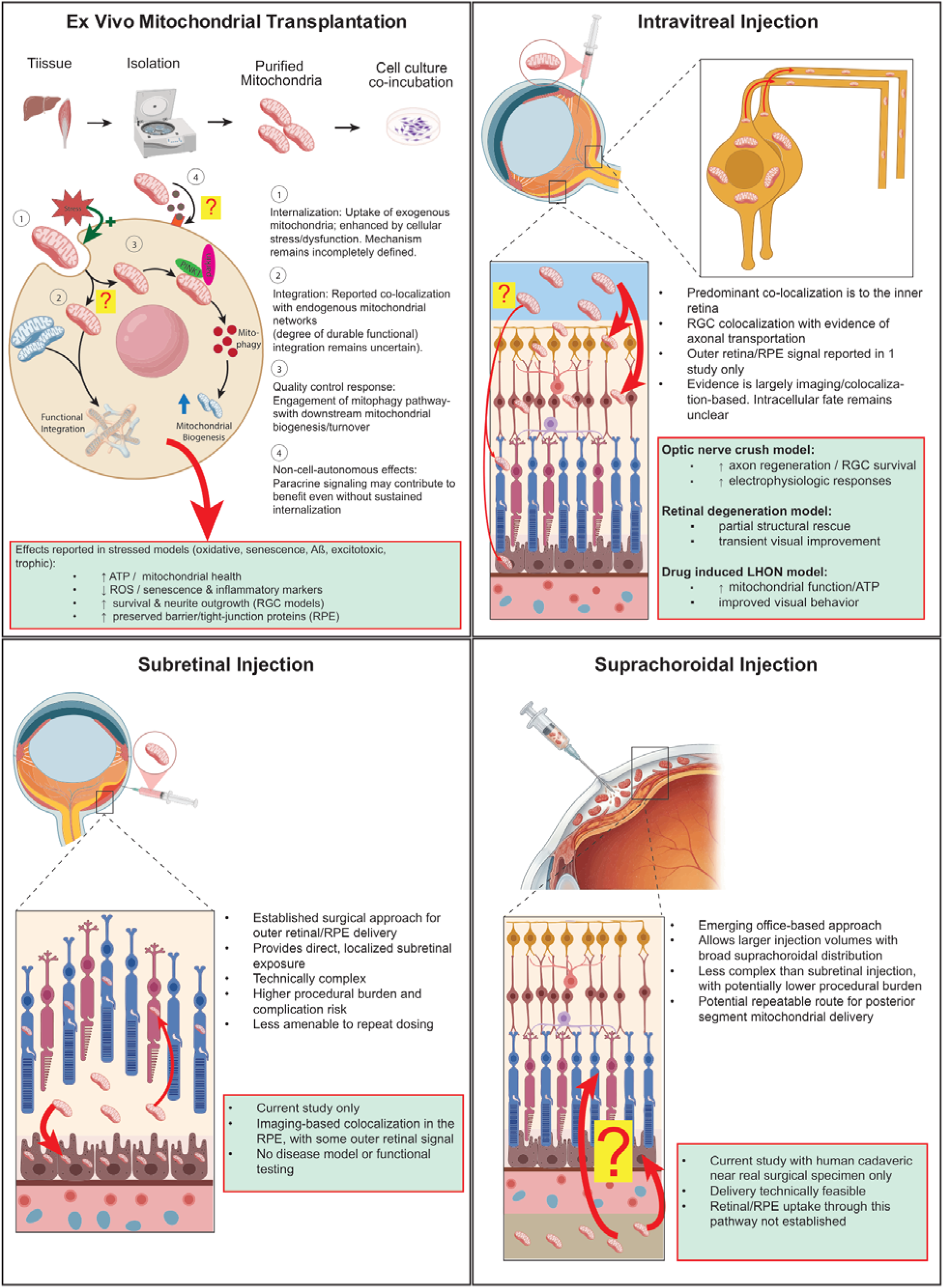
Ocular delivery routes for mitochondrial transplantation. Schematic of ex vivo transplantation, intravitreal injection, subretinal injection, and suprachoroidal injection. Reported localization differs by route, with uncertainty regarding mitochondrial uptake, retinal transport, intracellular fate, and therapeutic mechanism.

##### In Vitro Evidence

Aharoni-Simon and colleagues were among the first to demonstrate mitochondrial uptake in a retinal cell line.^80^ Mouse liver-derived mitochondria were co-incubated with the 661W cell line (an immortalized photoreceptor like cell line) and uptake was assessed using a strain-specific mtDNA qPCR method. Uptake was detectable within a few hours, peaking around 6 hours, and declining by 24 hours. Notably, hydrogen peroxide exposure increased uptake and persistence, consistent with prior reports that cellular stress can enhance mitochondrial internalization. The study used the tumor-derived 661W cell line, which captures only certain features of native photoreceptors, making in vivo translation uncertain. The use of mtDNA qPCR serves as an indirect marker and doesn’t necessarily confirm bioenergetically active organelles. In addition, mtDNA qPCR is an indirect readout and does not establish whether internalized mitochondria are intact or bioenergetically functional, and the study did not report functional outcomes.

Noh et al. ^81^ investigated mitochondrial transfer to stressed RPE cells using ARPE-19 cultures. They delivered stem cell-derived mitochondria by co-incubation and assessed uptake by MitoTracker labeling and confocal imaging. In models of replicative and oxidative senescence, umbilical cord-derived MSC mitochondria reduced senescence and oxidative stress markers and partially normalized mitochondrial quality-control pathways. Inflammatory markers also declined, suggesting an overall shift toward a more functional state. It is important to note however that these experiments were performed in ARPE-19 cells rather than primary human RPE limiting its interpretation. In another more AMD-relevant model, Noh and collogues ^82^ applied bone marrow-derived mitochondria to ARPE-19 cells exposed to oligomeric amyloid-β and reported improved metabolic function and preservation of barrier-associated proteins such as tighter junctional organization.

In a recent preprint, Ashok et al.^83^ evaluated mitochondrial transplantation across different neuronal cell lines in vitro relevant to retinal ganglion cell stress and survival. Exogenous mitochondria were labeled with MitoTracker Red and transferred to differentiated SH-SY5Y cells exposed to glutamate excitotoxicity and to PC12 cells under trophic deprivation (low NGF). In both settings, transplanted mitochondria were reported to colocalize with an endogenous mitochondrial signal and were associated with improved survival and neurite outgrowth, including increased neurite length and number as well as increased expression of neurite-associated markers. Mechanistically, the authors reported restoration of mitochondrial membrane potential, reduced reactive oxygen species, and increased ATP production. Importantly, they also extended these observations to primary mouse RGC cultures (isolated from P3 retinas), where exogenous mitochondria were associated with enhanced neurite outgrowth.

Together, these in vitro studies support feasibility of mitochondrial transfer to several retinal cell types and potential stress induced benefits but require confirmation in primary human cells, stronger evidence of functional donor-mitochondria contribution, and validation in in vivo settings.

##### In Vivo Evidence

A number of studies have evaluated mitochondrial transfer in vivo using mouse and rat models via the intravitreal delivery route. In an optic nerve crush model, Nascimento-dos-Santos et al^84^, reported that intravitreal injected MitoTracker-labeled donor mitochondria colocalized with TUJ1⁺ retinal ganglion cells in a tubular pattern, consistent with uptake and incorporation into a functional mitochondrial network. This was associated with improved bioenergetics and function with an improved ERG response and axonal growth. When repeated with lysed dysfunctional mitochondria this effect was not seen, underscoring the importance of mitochondrial integrity for the functional benefit. Interestingly, while lysed mitochondria appeared to boost early axon growth, only intact mitochondria supported a more sustained regeneration.

Wu et al. ^85^ tested mitochondrial transplantation via intravitreal delivery in RCS rats, a model of photoreceptor degeneration. They observed partial structural rescue and reduced retinal thinning on OCT imaging, as well as improved visual signal transmission early on. The effects were modest and diminished over time, and direct in vivo confirmation of mitochondrial integration was not attempted. So, it remains unclear whether this observed effect was due to direct mitochondrial integration or some other signaling mechanism.

Noh et al. ^82^ evaluated intravitreal delivery of human bone marrow-derived mitochondria in an in vivo mouse model following subretinal FITC-oAβ exposure, completing their earlier in vitro model. Donor mitochondria were detected within 24 hours on retinal cryosections and flat mounts using immunohistochemistry with a human-specific anti-mitochondria antibody. Interestingly, donor mitochondria were reported across all retinal layers, including the RPE, which contrast other studies including Nascimento-dos-Santos et al.,^84^ Wang et al.,^86^ and Ashok et al.,^83^ where intravitreal delivery resulted in a limited inner retinal localization. Functionally, mitochondrial transplantation was associated with a reduced FITC–oAβ signal and better preservation of RPE barrier architecture, including greater ZO-1 continuity compared with controls.

In another study, Wang et al. ^86^ developed a novel nanoengineered delivery platform by attaching PARKIN mRNA-loaded nanoparticles to isolated mitochondria, aiming to enhance uptake and mitophagy activity. In a rotenone-induced Leber’s Hereditary Optic Neuropathy (LHON) model, intravitreal delivery led to an increase PARKIN activity and improved mitochondrial function as well as a modest preservation of retinal structure and visual function. Surprisingly immunohistochemistry revealed only relatively sparse mitochondrial signal in the inner retina, raising the question of whether the observed benefits may reflect downstream PINK1-Parkin dependent mitophagy effects rather than functional integration of the transplanted mitochondria. An effect previously reported in a different setting.^52^ While the results are encouraging, the rotenone-induced model is an acute mitochondrial complex I inhibition rather than a true genetic LHON genotype, and whether similar rescue would occur in inherited LHON models remains to be seen.

Lastly, Ashok et al.^83^ (pre-print) also assessed the effect of in vivo mitochondrial transplantation in an optic nerve crush model. Donor mitochondria were isolated and labeled with MitoTracker Red. Prior to intravitreal delivery mitochondrial preparations underwent a quality-control step using membrane potential assays (JC-1) to ensure functionality. The authors reported mitochondrial signal colocalization within RBPM+ RGCs and inner retinal layers, consistent with other reports. Functionally, mitochondrial transplantation lead to improved RGC survival and improved electrophysiologic responses compared to controls. Anatomically they reported increased axon regeneration beyond the crush site using anterograde tracing. Interestingly MitoTracker Red positive mitochondria were identified along the regenerated axon nerve fibers, suggesting anterograde transport of mitochondria.

#### Current Challenges in Mitochondrial Delivery

Safety is a key translational concern for ocular mitochondrial transplantation since intraocular inflammation can lead to permanent vision loss with severe functional consequences to the patient. Although intravitreal and subretinal injections are now commonly performed in clinical settings, experience with other intraocular agents have highlighted the potential for rare but severe immune-mediated complications such as retinal vasculitis and vascular occlusion.^87–89^ Together, these events emphasize that emerging subcellular therapeutics such as mitochondrial transplantation must be evaluated not only for efficacy but also for inflammatory and vaso-occlusive risk.

The concern for potential immunogenicity of mitochondria is not unfounded. Mitochondria retain features of bacterial ancestry, including CpG-rich mitochondrial DNA and formylated peptides which have the potential to engage innate immune pathways such as TLR9 and formyl peptide receptors under certain contexts.^90,91^ At the same time, observations from the broader mitochondrial-transfer literature support cautious optimism that cell-free mitochondria, particularly autologous preparations, may be tolerated. Early clinical experience with autologous mitochondrial administration in non-ocular settings has not identified clear short-term safety signals.^92^ In addition, mitochondria can be detected in the circulation under physiologic conditions,^42^ supporting the idea that the body has existing mechanisms to accommodate extracellular mitochondrial material. Lastly, blood products with increased extracellular mitochondrial content are commonly administered, suggesting that exposure to extracellular mitochondrial material can be compatible with clinical use.^11^

Immunologic risk may depend on variables such as mitochondrial source (autologous vs heterologous), preparation purity (contaminating proteins, membrane fragments, free mtDNA), dose and repeat exposure, and the degree of mitochondrial damage (which could increase DAMP-like signaling). These considerations are particularly relevant for engineered mitochondrial platforms designed to enhance uptake or persistence.

For ocular applications specifically, safety data remains limited. In the nanoengineered mitochondrial model described by Wang et al. ^86^, the authors reported therapeutic benefit in an LHON model and did not emphasize overt inflammatory toxicity in their short-term assessments. However, existing studies were not powered or designed to detect rare inflammatory complications, and follow-up is often insufficient to exclude delayed immune events. Future studies should therefore incorporate dedicated safety endpoints including multimodal imaging to detect retinal vasculitis/nonperfusion, ocular cytokine profiling, and histology for microglial activation and vascular injury. Such evaluations will be essential for defining a therapeutic window in which mitochondrial delivery provides bioenergetic benefit without provoking vision-threatening inflammatory complications.

These uncertainties highlight that successful translation will require delivery methods that are not only effective, but also predictable and reproducible with respect to localization. Although there have been several previous studies indicating the feasibility of mitochondrial transplantation in the eye, there has been very little exploration as to the clinically relevant delivery routes of donor mitochondria into the eye. Here, we provide feasibility data using fluorescent reporter donor mitochondria to compare subretinal and intravitreal delivery in mice and to evaluate the procedural feasibility of suprachoroidal administration in a human cadaveric near-real surgical specimen, focusing on route-dependent distribution rather than therapeutic outcomes. (**Table 3**)

### Results - Feasibility of Ocular Mitochondrial Transplantation Across Clinically Relevant Delivery Routes

We utilized donor mitochondria with a genetically encoded DsRed reporter and assessed their distribution after three clinically relevant delivery routes: subretinal injection in postnatal mice, intravitreal injection in adult mice (including optic nerve head-directed delivery), and suprachoroidal administration in a near–real human surgical specimen model. In murine eyes, the DsRed_⁺_ signal was evaluated at early time points (6-48 hours) and at 7 days to capture initial localization and short-term persistence.

#### In vitro uptake of exogenous mitochondria by RPE cells

In vitro co-incubation with immortalized mouse RPE cells demonstrated a clear dose-dependent increase in intracellular DsRed_⁺_ mitochondria (**Figure 2**). To explore whether RPE cells can internalize exogenous mitochondria in principle, we performed a complementary in vitro assay using RPE cells. Confocal microscopy confirmed robust mitochondrial uptake in a dose dependent manner, supporting the plausibility that mitochondria could be internalized if they reached the RPE from a suprachoroidal injection in vivo. Isolated DsRed_⁺_ mitochondria were incubated with retinal pigment epithelial (RPE) cells in culture for 18 hours (**Figure 2A**). Confocal microscopy demonstrated DsRed_⁺_ signal co-localized with the RPE cells, indicating possible mitochondrial uptake. The number of internalized mitochondria increased in a dose dependent manner (**Figure 2B**).

**Figure 2.**
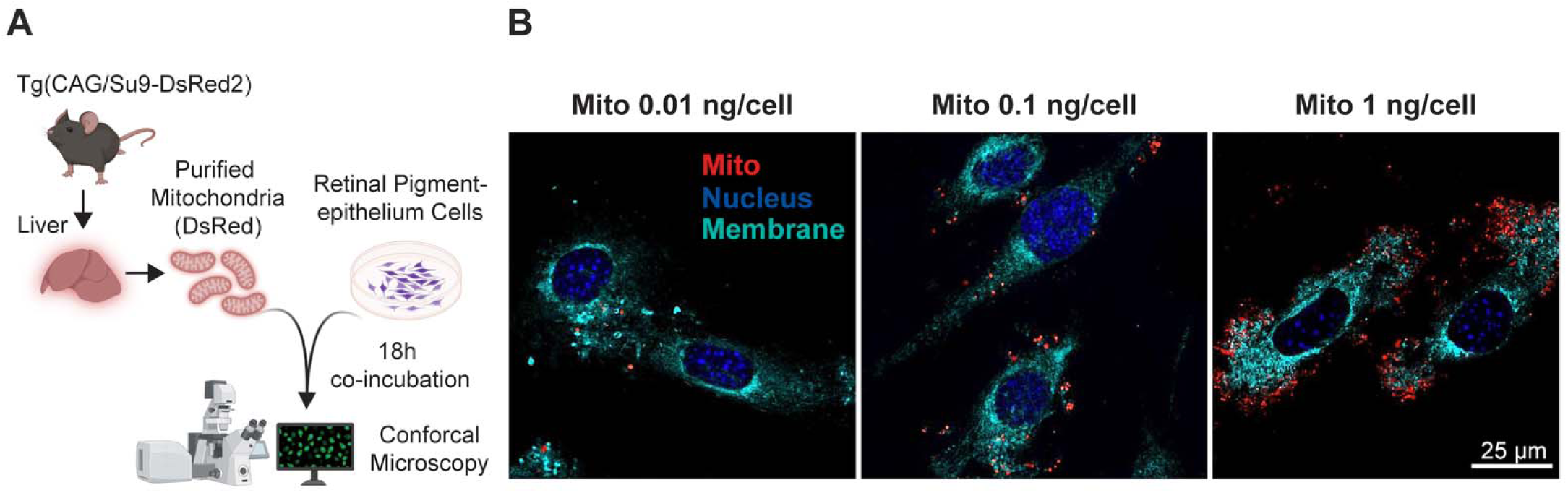
Dose-dependent uptake of DsRed^⁺^ mitochondria by cultured RPE cells. **(A)** Schematic of the in vitro uptake assay. DsRed^⁺^ mitochondria were purified from liver of Tg(CAG/Su9-DsRed2) donor mice and co-incubated with cultured retinal pigment epithelium (RPE) cells for 18 hours, followed by confocal microscopy.**(B)** Representative confocal images demonstrate a clear dose-dependent increase in DsRed^⁺^ mitochondrial signal within RPE cells following exposure to 0.01, 0.1, or 1 ng mitochondria per cell. DsRed^⁺^ mitochondria (red) appear as punctate cytoplasmic signal with prominent perinuclear distribution. Nuclei are labeled in blue and plasma membrane in cyan. Scale bar, 25 µm

#### Subretinal injection of Isolated Mitochondria

Subretinal injections of DsRed+ mitochondria were performed in 7-day old mice pups. Mouse eyes were harvested 6, 24, 48 hours and 7 days after injection in the subretinal space (**Figure 3A**). This resulted in a robust fluorescence signal within the subretinal space 6h after injection (**Figure 3B-C**). Higher magnification showed DsRed+ dot-like fluorescence co-localizing with the retinal pigmented epithelium (RPE) at 6 hours post-injection (**Figure 3D-E**). Similar staining pattern was seen at 24h. By post-injection day 7 the DsRed+ signal with dot-like pattern was no longer detectable. Subretinal delivery produced the most robust and anatomically specific mitochondrial colocalization in vivo with a prominent DsRed_⁺_ signal aligning with the RPE layer at both 6 and 24 hours

**Figure 3.**
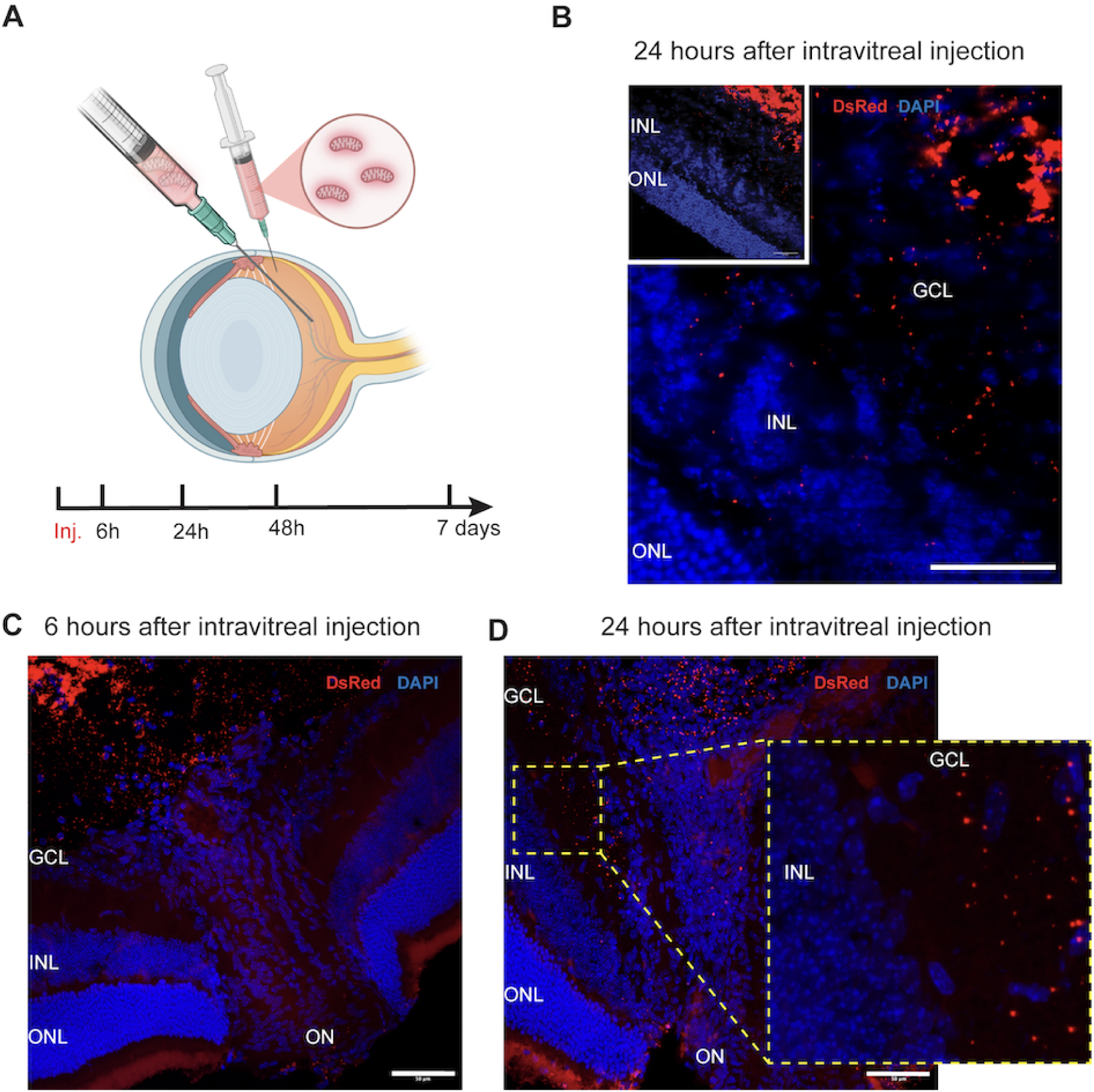
Subretinal mitochondrial delivery. **(A)** Schematic representation of the experimental design. Isolated DsRed_⁺_ mitochondria were injected into the subretinal space of P7 mouse pups. Eyes were collected for immunohistochemistry at 6 hours, 24 hours, 48 hours, and 7 days post-injection. **(B)** Retinal eyecup 6 hours after subretinal injection showing DsRed_⁺_ fluorescence, highlighting subretinal distribution of transplanted mitochondria. **(C)** Retinal cross-section 24 hours post-subretinal injection in adult mice, demonstrating DsRed_⁺_ signal in the subretinal space. Scale bar: 500 µm. **(E)** Higher magnification of retinal cross-section 24 hours post-injection showing DsRed_⁺_ mitochondria colocalizing with retinal pigment epithelium (RPE) and the outer nuclear layer (ONL). Scale bar: 20 µm. *Abbreviations: ONL= outer nuclear layer, INL= inner nuclear layer, GC= ganglion cell layer*

#### Intravitreal injection of Isolated Mitochondria

Intravitreal injection of DsRed+ mitochondria into 5-week-old adult mice in the mid cavity and directed towards the optic nerve utilizing a longer needle (**Figure 3A**) resulted in a strong DsRed+ fluorescence signal within the vitreous cavity. 24-hours post-injection (**Figure 3B**) shows scattered dot-like DsRed+ fluorescence in the ganglion cell layer (GCL), inner plexiform layer (IPL) and inner nuclear layer (INL), with notable signal intensity reductions by 48 hours and was greatly diminished on post injection day 7. After targeted intravitreal injection towards the optic nerve head, DsRed+ mitochondria were more densely observed within the optic nerve head ganglion cell fibers at 6-or 24-hours post-injection (**Figure 4C-D**). The DsRed+ mitochondria appeared to migrate more distally along the optic nerves over time (**Figure 4D magnification)**. Like the subretinal injection, the fluorescence signal nearly resolved by post-injection day 7.

**Figure 4.**
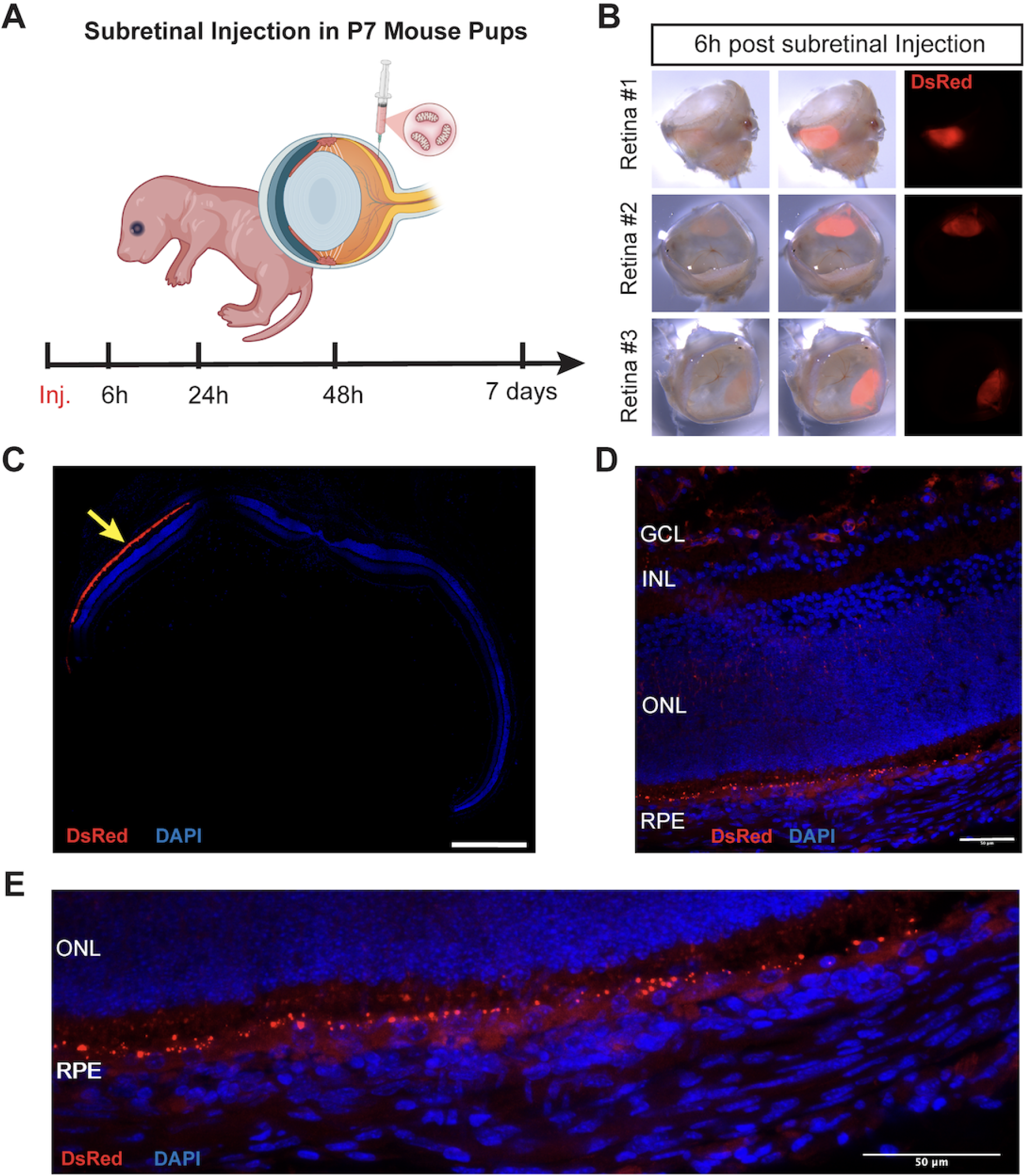
Intravitreal mitochondrial delivery. **(A)** Schematic representation of the experimental design. Isolated DsRed_⁺_ mitochondria were injected into the mid-vitreous cavity of adult (5-week-old) CD1 mice. Using a longer needle, mitochondria were also delivered closer to the optic nerve region. **(B)** Retinal cross-section 24 hours post-intravitreal injection. Higher magnification of the highlighted region (white box; top left) shows DsRed_⁺_ mitochondria localized to the ganglion cell layer (GCL), inner plexiform layer (IPL), and inner nuclear layer (INL). Scale bar: 50µm. **(C)** Retinal cross-section through the optic nerve head at 6 hours after targeted injection towards the optic nerve head shows DsRed_⁺_ mitochondria predominantly localized to retinal ganglion cell axons at the proximal optic nerve head. Scale bar: 50µm **(D)** Retinal cross-section through the optic nerve head at 24 hours after targeted injection towards the optic nerve head reveals DsRed_⁺_ mitochondria in more distal segments of retinal ganglion cell axons (highlighted in the magnified box). DAPI stains nuclei; DsRed labels transplanted mitochondria. Scale bar: 50µm *Abbreviations: ONL= outer nuclear layer, INL= inner nuclear layer, GC= ganglion cell layer, ON= optic nerve*

#### Suprachoroidal Injection of Isolated Mitochondria

To test the feasibility of injecting mitochondria in the human suprachoroidal space, we used a cadaveric human near-real surgical specimen and the novel Everads suprachoroidal injection device (**Figure 5**). We performed 8 injections in three near-real surgical specimens. The procedure demonstrated consistent access to the suprachoroidal space without instrument clogging or backflush of the mitochondrial suspension. Visual confirmation of successful delivery was achieved using brilliant blue dye, which showed appropriate distribution within the suprachoroidal space. The mitochondrial suspension maintained its integrity throughout the injection process, with no visible aggregation or precipitation that would compromise delivery. Cutdown of the sclera with an MVR blade after injection resulted in immediate release of injectate from the incision, further confirming suprachoroidal space placement. Post-injection examination revealed no evidence of scleral perforation or unintended penetration into the vitreous cavity, supporting the controlled nature of the delivery technique.

**Figure 5.**
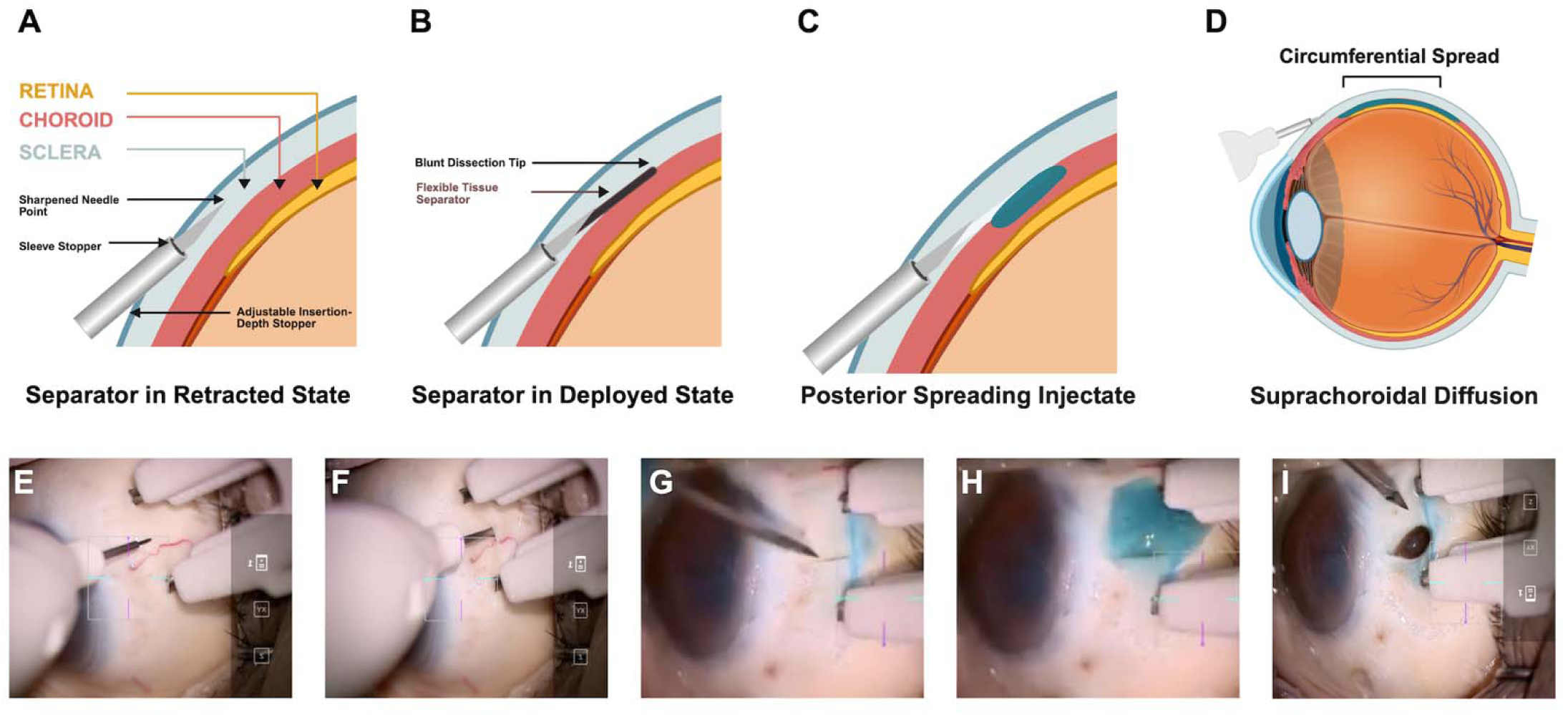
Suprachoroidal mitochondria delivery. **(A–D)** Schematic illustration of the suprachoroidal delivery device (Everads Therapy Ltd), which incorporates a non-sharp nitinol tissue separator for tangential blunt dissection through the sclera to access the suprachoroidal space while maintaining choroidal integrity. **(A)** Tangential scleral insertion with the separator in the retracted state and insertion depth controlled by an adjustable stopper. **(B)** Deployment of the blunt tissue separator to establish a dissection plane toward the suprachoroidal space. **(C)** Posterior spread of injectate within the suprachoroidal space. **(D)** Circumferential diffusion of injectate throughout the suprachoroidal space across the posterior segment. **(E–I)** Representative procedural images showing device insertion and suprachoroidal delivery. **(E, F)** The eye was stabilized, the device positioned, and the blue-stained solution was injected without any reflux. **(G)** To confirm suprachoroidal injection, an MVR blade was used to perform a scleral cut down. (**H**) There was egress of the blue-stained injectate. (**I**) Further exposure of the suprachoroidal space showed the choroid remained intact, consistent with successful suprachoroidal delivery and no overt choroidal penetration or disruption.

## Discussion: Translational Considerations for Mitochondrial Transplantation in the Eye

Mitochondrial dysfunction is increasingly recognized as a key contributor to major blinding disorders, reflecting the high energetic demands and vulnerability to energy disruption of the neural retina and retinal pigment epithelium. As the population ages, age-related conditions such as AMD and glaucoma are projected to rise significantly, reaching 288 million and 110 million cases by 2040, respectively, highlighting the need for innovative treatment options. The growing mitochondrial transplantation literature shows improvement bioenergetic function and promotion of tissue-protective effects in various disease models, which inspired interest in mitochondrial transplantation in the eye. Although the literature on ocular mitochondrial transplantation is relatively sparse, several in vivo studies have shown potential for beneficial effects. For instance, in 2 acute optic nerve injury models, intravitreal mitochondrial delivery resulted in improved bioenergetic function and enhanced axon regeneration.^83,84^ Furthermore, in a retinal degeneration model mitochondrial transplantation showed a modest structural and functional rescue.^85^ In a RPE injury context, delivery of exogenous mitochondria improved RPE architecture and barrier-related features.^82^ Lastly, in a drug-induced LHON-relevant model, an engineered intravitreal mitochondrial delivery approach restored complex I activity and ATP production, resulting in improved visual behavior and highlighting potential relevance for primary mitochondrial diseases.^86^ These studies indicate the need for further investigation while also emphasizing that the field has not yet reached a consensus on the mechanisms or optimal delivery methods.

Over the past several decades, intraocular drug delivery techniques have evolved substantially. The widespread adoption of intravitreal injection of anti-VEGF agents has revolutionized the treatment of posterior segment diseases by providing scalable and safe office-based treatments.^93^ More recently, subretinal injections have expanded therapeutic possibilities by enabling direct access to the outer retina and RPE allowing the effective delivery of gene therapy.^94^ In parallel, suprachoroidal injection has emerged as an alternative strategy, providing a similar exposure to the posterior segment, while avoiding a more complex subretinal delivery in the operating room and its associated complications.^93^ These recent advancements in surgical delivery provide a pathway for the development of ocular mitochondrial transplantation techniques.

In contrast to many other tissues, the eye is a highly structured organ with distinct anatomical compartments that limit where material can be distributed. As a result, the delivery route is a key factor in the efficacy of transplanted mitochondria in accessing the cell type of interest, partly due to route-specific barriers to tissue penetration. Procedural complexity further influences the practicality of repeat dosing. In this context, our feasibility data supplements the existing literature by directly comparing route-dependent donor mitochondrial localization using a genetically encoded fluorescent mitochondrial reporter mouse (DsRed_⁺_ mitochondria). This approach enables direct tracking of the donor-derived signal without reliance on dye-based labeling such as MitoTracker, which can be confounded by dye transfer.

In murine experiments, we observed delivery-dependent patterns of the DsRed_⁺_ signal. Intravitreal delivery produced punctate signal predominantly within inner retinal layers, with enrichment near the optic nerve head when injections were directed toward this region. This inner retina predominant distribution is consistent with multiple prior intravitreal studies reporting localization within the RGC and deeper inner retinal layers.^83,84,86^ Notably, we did not observe outer retinal or RPE localization after intravitreal delivery, in contrast to reports relying on antibody-based detection,^82^ which may be influenced by differences in injury context, tissue permeability, or detection methodology. In contrast, subretinal delivery yielded a prominent signal at the outer retina and RPE interface, supporting subretinal injection as an anatomically direct strategy for RPE-adjacent exposure. This is further supported by our own and others RPE cell culture experiments showing colocalization within cell boundaries after exogenous mitochondria exposure.^81,82^ Although subretinal delivery may be attractive for outer retinal targets, it will be important to determine whether a short-lived exposure of mitochondria will be sufficient to induce a clinically durable effect

Overall, our results and the existing literature indicate that intravitreal injection is a feasible method for mitochondrial delivery to the inner retina. Our data also highlights subretinal delivery as a potentially compelling strategy to deliver mitochondria to the outer retina and RPE. However, it is important to keep in mind that subretinal injections are procedurally complex with a higher risk profile, and, therefore, may be less suitable for repeated administrations. These limitations motivate continued exploration of less invasive alternatives, including suprachoroidal delivery and engineered approaches aimed at improving tissue penetration to reach outer retinal targets via intravitreal delivery.

Suprachoroidal injection has recently expanded as an office-based approach using larger injectable volumes that can distribute broadly within the suprachoroidal space; however, experience has largely been limited to small molecules and gene-therapy platforms. The Everads Injector utilizes a 30-gauge needle with a non-sharp nitinol tissue separator that creates access to the suprachoroidal space via tangential blunt dissection. Studies show this injector is tolerable and effective in humans,^95^ and in non-human primate studies, to enable rapid delivery of injectate to the posterior segment and macula. ^95–97^ In that regard, our near-real human surgical specimen experiments establish the procedural feasibility of delivering a mitochondrial suspension to the suprachoroidal space using a dedicated Everads injector; nevertheless, whether suprachoroidal delivered mitochondria reliably access the neural retina or RPE in vivo remains an open and critical question.

It is important to note limitations that apply across all studies, including our own. Most studies rely primarily on confocal imaging-based localization which is valuable for assessing colocalization but cannot distinguish true intracellular uptake from extracellular association. This lack of subcellular resolution limits the mechanistic interpretation. It remains unclear whether the effects seen reflect functional integration of donor mitochondria into existing mitochondrial networks, PINK1-PARKIN-mediated mitophagy downstream signaling effects, or an independent paracrine effect unrelated to mitochondrial incorporation. Higher-resolution approaches, such as electron microscopy with mitochondrial labeling (e.g APEX2), are needed to resolve intracellular localization and track the subcellular fate. In addition to high-resolution imaging, more granular mechanistic investigations using a multimodal approach, such as mtDNA tracking and targeted disruption of individual pathways like mitophagy, will be necessary to distinguish the dominant mechanisms underlying the reported functional outcomes.

Another important translational gap is that ocular mitochondrial transplantation has not yet been tested in the most clinically relevant chronic disease models such as age-related macular degeneration, glaucoma, and diabetic retinopathy where mitochondrial dysfunction plays a central role. In these chronic disease settings, it remains to be seen whether a single transplantation can result in a durable functional benefit or whether repeated administrations would be necessary to achieve and sustain a clinically meaningful effect.

In summary, the available literature and our data support the feasibility of ocular mitochondrial transplantation and validates continued investigation. However, major uncertainties regarding optimal delivery, mechanism of action, persistence, and safety remain. Our feasibility data on mitochondria delivery add an anatomic perspective to a heterogeneous literature and underscores the importance of selecting an appropriate delivery rout to reach the cells of interest. Future studies with more detailed mechanistic work, including donor mtDNA quantification, ultrastructural localization with determination of subcellular fate, longitudinal functional testing and pathway modulation in disease-relevant models, will be needed to clarify mechanism, guide dosing strategies, and determine whether mitochondria transplantation can deliver a durable benefit in the eye.

## Methods

### Narrative review

We conducted a focused, non-systematic narrative review of mitochondrial transplantation with an emphasis on ocular applications and posterior-segment delivery paradigms. The goal of the review was to summarize (1) the biologic rationale for mitochondrial transplantation, (2) mechanisms of intercellular mitochondrial transfer, (3) ocular disease contexts in which mitochondrial dysfunction is implicated, and (4) preclinical in vitro and in vivo studies that have evaluated mitochondrial transplantation in ocular tissues. Studies were identified through targeted searches of PubMed and Google Scholar using combinations of the terms “mitochondrial transplantation”, “mitochondrial transfer”, “retina,” “retinal ganglion cell”, “retinal pigment epithelium”, “intravitreal”, “subretinal”, and “suprachoroidal” and through citation tracking of relevant primary manuscripts and reviews. Because this was a narrative review, we did not predefine inclusion and exclusion criteria, perform a formal risk-of-bias assessment, or apply PRISMA methodology. Findings from the ocular literature were synthesized qualitatively and organized by experimental context (in vitro vs. in vivo) and delivery route.

### Animals

The experiments were conducted in accordance with the ARVO Statement for the Use of Animals in Ophthalmic and Vision Research and approved by Stanford’s Institutional Animal Care and Use Committee. For intravitreal and subretinal mitochondria injections we utilized CD-1 mice obtained from Charles River.

### Mitochondria Isolation

Mitochondria were isolated from C57Bl6/J mouse livers expressing red fluorescent protein fused to a mitochondrial import sequence (mitochondria-DsRed) using a previously published protocol.^98,99^ Briefly, following cervical dislocation, livers were excised, rinsed, and homogenized in ice-cold mitochondrial isolation buffer. Cell debris and nuclei were removed by low-speed centrifugation, followed by successive high-speed spins to eliminate contaminants. The pure mitochondria were isolated from crude mitochondrial fraction by Percoll gradient ultracentrifugation, and the mitochondrial band was collected in PBS.^100^

### Mitochondrial transplantation

CD-1 mice (Charles River) were used for all experiments and anesthetized with ketamine and xylazine administered via intraperitoneal injection. Once appropriate anesthesia was achieved, the mice were positioned under an operating microscope for mitochondrial transplantation procedures.

### Subretinal Injection

Subretinal injections were performed in postnatal day 7 mouse pups as previously described.^101^ Briefly, a small scleral incision was made at or just posterior to the globe’s equator by gently scratching the surface with a 15-degree microsurgery blade to access the subretinal space. Using a pre-prepared microinjection needle, pulled from glass capillary tubing, 1 µL of the of DsRed+ Mitochondria solution was injected into the subretinal space using a microinjection system (Femtojet Eppendorf).

### Intravitreal Injection

Intravitreal injections were performed in 5-week-old CD1 mice. The procedure was similarl to the subretinal injection, with the key distinction being that the incision was extended until clear vitreous was encountered. The solution containing the of DsRed+ Mitochondria was then injected into the vitreous cavity. To achieve a higher concentration of mitochondria near the optic nerve head, a longer needle was utilized and directed toward the optic nerve head instead of aiming toward the center of the vitreous cavity.

### Suprachoroidal Injection

Cadaveric human near-real surgical specimens were obtained from consented donors within 24 hours post-mortem and maintained at 4°C until use. The Everads suprachoroidal injection system was used to deliver mitochondria to the suprachoroidal space. The device features a 30-gauge needle with a non-sharp nitinol tissue separator that creates a controlled pathway from the sclera to the choroid through tangential blunt dissection. Following sterile preparation, the injector was positioned tangentially to the scleral surface approximately 3-4 mm posterior to the limbus. The tissue separator was deployed to create access to the suprachoroidal space, then retracted to leave an open channel. A 150 μL solution containing mitochondria mixed with brilliant blue dye (for visualization) was then injected through the established pathway. The injection volume was delivered at a controlled rate to ensure proper distribution within the suprachoroidal space.

### Immunofluorescence

Murine eyes were enucleated at 6, 24, and 48 hours, as well as 7 days following mitochondrial transplantation, and immediately fixed in 4% paraformaldehyde for 2 hours at room temperature. After fixation, the eyes were rinsed with PBS, and treated with sucrose gradient (5%, 15% and 30% sucrose in PBS), immersed in OCT, and frozen and stored at -80. Frozen eyes were cryo-sectioned into 20μm slices, rinsed with PBS, and blocked using PBS containing 1%BSA, 10% Donkey Serum and 0.1% TritonX100. Sections were subsequently stained with DAPI (ThermoFisher, D1306) for 15 minutes at room temperature.

### Retinal Pigment Epithelium Cell Culture

Glass slides were pre-coated with Applied Cell Extracellular Matrix (ABM, Cat# G422). Immortalized mouse RPE cells (ABM, Cat# T05740) were cultured in Prigrow III Medium (ABM, Cat# TM003) supplemented with 10% non-heat-inactivated FBS and 1× Penicillin/Streptomycin.

For mitochondrial transfer experiments, DsRed-labeled mitochondria were added to 40,000 RPE cells at the amounts indicated in the corresponding figure legends (µg). The highest concentration corresponded to approximately 1 ng/cell, with serial 10-fold dilutions used for lower concentrations, performed overnight.

After incubation, wells were washed, and plasma membranes were stained with CellMask Deep Red (5 min) followed by three washes. Cells were then fixed with 2% paraformaldehyde (PFA) for 20 min at room temperature and counterstained with Hoechst (5 µg/mL, 5 min). Mounting was performed using Dako mounting medium.

## Acknowledgments

E.B. is recipient of a Merit Award from the Fonds de Recherche en Santé du Québec. We thank MaryAnn Mahajan for editorial assistance.

## Author Contributions Statement

**–** Dr. Mahajan had full access to all the data in the study and took responsibility for the integrity of the data and the accuracy of the data analysis. Study concept and design: BC, VBM, SW. Acquisition of data: BC, TCY, CHL, MRW, EB, MP, YB, SP, SW, VBM. Analysis and interpretation of data: BC, TCY, CHL, MRW, EB, AS, MP, YB, SP, TB, DRPA, SW, VBM. Drafting of the manuscript: BC, TCY, SW, VBM. Critical revision of the manuscript for important intellectual content: all authors. Obtained funding: SW, VBM. Administrative, technical, and material support: SW, VBM. Study supervision: VBM.

## Institutional Review Board Statement

**–** The study was approved by the Stanford University Institutional Review Board and adhered to the tenets set forth in the Declaration of Helsinki.

## Financial Support

**–** TCY is supported by NIH grant 2T32EY027816. VBM is supported by (R01EY037830, R01EY030151, and P30EY026877), Research to Prevent Blindness, New York, New York, the and Alan and Irene Adler Ocular Oncology Center at Stanford.

## Disclosures

EB and VBM are advisors for MitrixBio

EB holds patents related to the detection of extracellular mitochondria

## Notes

### Competing Interest Statement

The authors have declared no competing interest.

## References

1. van der Bliek AM, Sedensky MM, Morgan PG. Cell Biology of the Mitochondrion. Genetics. 2017;207(3):843–871. doi:10.1534/genetics.117.300262

2. Labbé K, Murley A, Nunnari J. Determinants and functions of mitochondrial behavior. Annu Rev Cell Dev Biol. 2014;30:357–391. doi:10.1146/annurev-cellbio-101011-155756

3. Pfanner N, Warscheid B, Wiedemann N. Mitochondrial proteins: from biogenesis to functional networks. Nat Rev Mol Cell Biol. 2019;20(5):267–284. doi:10.1038/s41580-018-0092-0

4. Wong-Riley MTT. Energy metabolism of the visual system. Eye Brain. 2010;2:99–116. doi:10.2147/EB.S9078

5. Liu L, Trimarchi JR, Smith PJS, Keefe DL. Mitochondrial dysfunction leads to telomere attrition and genomic instability. Aging Cell. 2002;1(1):40–46. doi:10.1046/j.1474-9728.2002.00004.x

6. Zhang ZQ, Xie Z, Chen SY, Zhang X. Mitochondrial dysfunction in glaucomatous degeneration. Int J Ophthalmol. 2023;16(5):811–823. doi:10.18240/ijo.2023.05.20

7. Somasundaram I, Jain SM, Blot-Chabaud M, et al. Mitochondrial dysfunction and its association with age-related disorders. Front Physiol. 2024;15:1384966. doi:10.3389/fphys.2024.1384966

8. Chen BS, Harvey JP, Gilhooley MJ, Jurkute N, Yu-Wai-Man P. Mitochondria and the eye—manifestations of mitochondrial diseases and their management. Eye. 2023;37(12):2416–2425. doi:10.1038/s41433-023-02523-x

9. Du Y, Veenstra A, Palczewski K, Kern TS. Photoreceptor cells are major contributors to diabetes-induced oxidative stress and local inflammation in the retina. Proc Natl Acad Sci U S A. 2013;110(41):16586–16591. doi:10.1073/pnas.1314575110

10. Oh JK, Lima de Carvalho JR, Nuzbrokh Y, et al. Retinal Manifestations of Mitochondrial Oxidative Phosphorylation Disorders. Invest Ophthalmol Vis Sci. 2020;61(12):12. doi:10.1167/iovs.61.12.12

11. Borcherding N, Brestoff JR. The power and potential of mitochondria transfer. Nature. 2023;623(7986):283–291. doi:10.1038/s41586-023-06537-z

12. Klopstock T, Yu-Wai-Man P, Dimitriadis K, et al. A randomized placebo-controlled trial of idebenone in Leber’s hereditary optic neuropathy. Brain. 2011;134(Pt 9):2677–2686. doi:10.1093/brain/awr170

13. Sadun AA, Chicani CF, Ross-Cisneros FN, et al. Effect of EPI-743 on the clinical course of the mitochondrial disease Leber hereditary optic neuropathy. Arch Neurol. 2012;69(3):331–338. doi:10.1001/archneurol.2011.2972

14. Mettu PS, Allingham MJ, Cousins SW. Phase 1 Clinical Trial of Elamipretide in Dry Age-Related Macular Degeneration and Noncentral Geographic Atrophy: ReCLAIM NCGA Study. Ophthalmol Sci. 2022;2(1):100086. doi:10.1016/j.xops.2021.100086

15. Age-Related Eye Disease Study 2 Research Group. Lutein + zeaxanthin and omega-3 fatty acids for age-related macular degeneration: the Age-Related Eye Disease Study 2 (AREDS2) randomized clinical trial. JAMA. 2013;309(19):2005–2015. doi:10.1001/jama.2013.4997

16. Williams PA, Harder JM, Cardozo BH, Foxworth NE, John SWM. Nicotinamide treatment robustly protects from inherited mouse glaucoma. Commun Integr Biol. 2018;11(1):e1356956. doi:10.1080/19420889.2017.1356956

17. Nashine S, Cohen P, Chwa M, et al. Humanin G (HNG) protects age-related macular degeneration (AMD) transmitochondrial ARPE-19 cybrids from mitochondrial and cellular damage. Cell Death Dis. 2017;8(7):e2951. doi:10.1038/cddis.2017.348

18. Feuer WJ, Schiffman JC, Davis JL, et al. Gene Therapy for Leber Hereditary Optic Neuropathy: Initial Results. Ophthalmology. 2016;123(3):558–570. doi:10.1016/j.ophtha.2015.10.025

19. Ji MH, Kreymerman A, Belle K, et al. The Present and Future of Mitochondrial-Based Therapeutics for Eye Disease. Transl Vis Sci Technol. 2021;10(8):4. doi:10.1167/tvst.10.8.4

20. Roger AJ, Muñoz-Gómez SA, Kamikawa R. The Origin and Diversification of Mitochondria. Curr Biol. 2017;27(21):R1177–R1192. doi:10.1016/j.cub.2017.09.015

21. Spees JL, Olson SD, Whitney MJ, Prockop DJ. Mitochondrial transfer between cells can rescue aerobic respiration. Proc Natl Acad Sci U S A. 2006;103(5):1283–1288. doi:10.1073/pnas.0510511103

22. Ahmad T, Mukherjee S, Pattnaik B, et al. Miro1 regulates intercellular mitochondrial transport & enhances mesenchymal stem cell rescue efficacy. EMBO J. 2014;33(9):994–1010. doi:10.1002/embj.201386030

23. Plotnikov EY, Khryapenkova TG, Vasileva AK, et al. Cell-to-cell cross-talk between mesenchymal stem cells and cardiomyocytes in co-culture. J Cell Mol Med. 2008;12(5A):1622–1631. doi:10.1111/j.1582-4934.2007.00205.x

24. Liu K, Ji K, Guo L, et al. Mesenchymal stem cells rescue injured endothelial cells in an in vitro ischemia-reperfusion model via tunneling nanotube like structure-mediated mitochondrial transfer. Microvasc Res. 2014;92:10–18. doi:10.1016/j.mvr.2014.01.008

25. Chen J, Zhong J, Wang L lin, Chen Y ying. Mitochondrial Transfer in Cardiovascular Disease: From Mechanisms to Therapeutic Implications. Front Cardiovasc Med. 2021;8:771298. doi:10.3389/fcvm.2021.771298

26. Islam MN, Das SR, Emin MT, et al. Mitochondrial transfer from bone-marrow-derived stromal cells to pulmonary alveoli protects against acute lung injury. Nat Med. 2012;18(5):759–765. doi:10.1038/nm.2736

27. Sinclair KA, Yerkovich ST, Hopkins PMA, Chambers DC. Characterization of intercellular communication and mitochondrial donation by mesenchymal stromal cells derived from the human lung. Stem Cell Res Ther. 2016;7(1):91. doi:10.1186/s13287-016-0354-8

28. Jackson MV, Morrison TJ, Doherty DF, et al. Mitochondrial Transfer via Tunneling Nanotubes is an Important Mechanism by Which Mesenchymal Stem Cells Enhance Macrophage Phagocytosis in the In Vitro and In Vivo Models of ARDS. Stem Cells. 2016;34(8):2210–2223. doi:10.1002/stem.2372

29. Mistry JJ, Marlein CR, Moore JA, et al. ROS-mediated PI3K activation drives mitochondrial transfer from stromal cells to hematopoietic stem cells in response to infection. Proc Natl Acad Sci U S A. 2019;116(49):24610–24619. doi:10.1073/pnas.1913278116

30. Jiang D, Gao F, Zhang Y, et al. Mitochondrial transfer of mesenchymal stem cells effectively protects corneal epithelial cells from mitochondrial damage. Cell Death Dis. 2016;7(11):e2467. doi:10.1038/cddis.2016.358

31. Osswald M, Jung E, Sahm F, et al. Brain tumour cells interconnect to a functional and resistant network. Nature. 2015;528(7580):93–98. doi:10.1038/nature16071

32. Lu J, Zheng X, Li F, et al. Tunneling nanotubes promote intercellular mitochondria transfer followed by increased invasiveness in bladder cancer cells. Oncotarget. 2017;8(9):15539–15552. doi:10.18632/oncotarget.14695

33. Nitzan K, Benhamron S, Valitsky M, et al. Mitochondrial Transfer Ameliorates Cognitive Deficits, Neuronal Loss, and Gliosis in Alzheimer’s Disease Mice. J Alzheimers Dis. 2019;72(2):587–604. doi:10.3233/JAD-190853

34. Babenko VA, Silachev DN, Zorova LD, et al. Improving the Post-Stroke Therapeutic Potency of Mesenchymal Multipotent Stromal Cells by Cocultivation With Cortical Neurons: The Role of Crosstalk Between Cells. Stem Cells Transl Med. 2015;4(9):1011–1020. doi:10.5966/sctm.2015-0010

35. Boukelmoune N, Chiu GS, Kavelaars A, Heijnen CJ. Mitochondrial transfer from mesenchymal stem cells to neural stem cells protects against the neurotoxic effects of cisplatin. Acta Neuropathol Commun. 2018;6(1):139. doi:10.1186/s40478-018-0644-8

36. Crewe C, Funcke JB, Li S, et al. Extracellular vesicle-based interorgan transport of mitochondria from energetically stressed adipocytes. Cell Metab. 2021;33(9):1853–1868.e11. doi:10.1016/j.cmet.2021.08.002

37. Rosina M, Ceci V, Turchi R, et al. Ejection of damaged mitochondria and their removal by macrophages ensure efficient thermogenesis in brown adipose tissue. Cell Metab. 2022;34(4):533–548.e12. doi:10.1016/j.cmet.2022.02.016

38. Nicolás-Ávila JA, Lechuga-Vieco AV, Esteban-Martínez L, et al. A Network of Macrophages Supports Mitochondrial Homeostasis in the Heart. Cell. 2020;183(1):94–109.e23. doi:10.1016/j.cell.2020.08.031

39. Boudreau LH, Duchez AC, Cloutier N, et al. Platelets release mitochondria serving as substrate for bactericidal group IIA-secreted phospholipase A2 to promote inflammation. Blood. 2014;124(14):2173–2183. doi:10.1182/blood-2014-05-573543

40. Hayakawa K, Esposito E, Wang X, et al. Transfer of mitochondria from astrocytes to neurons after stroke. Nature. 2016;535(7613):551–555. doi:10.1038/nature18928

41. Borcherding N, Jia W, Giwa R, et al. Dietary lipids inhibit mitochondria transfer to macrophages to divert adipocyte-derived mitochondria into the blood. Cell Metab. 2022;34(10):1499–1513.e8. doi:10.1016/j.cmet.2022.08.010

42. Al Amir Dache Z, Otandault A, Tanos R, et al. Blood contains circulating cell-free respiratory competent mitochondria. FASEB J. 2020;34(3):3616–3630. doi:10.1096/fj.201901917RR

43. Stephens OR, Grant D, Frimel M, et al. Characterization and origins of cell-free mitochondria in healthy murine and human blood. Mitochondrion. 2020;54:102–112. doi:10.1016/j.mito.2020.08.002

44. Joshi AU, Minhas PS, Liddelow SA, et al. Fragmented mitochondria released from microglia trigger A1 astrocytic response and propagate inflammatory neurodegeneration. Nat Neurosci. 2019;22(10):1635–1648. doi:10.1038/s41593-019-0486-0

45. Brestoff JR, Wilen CB, Moley JR, et al. Intercellular Mitochondria Transfer to Macrophages Regulates White Adipose Tissue Homeostasis and Is Impaired in Obesity. Cell Metab. 2021;33(2):270–282.e8. doi:10.1016/j.cmet.2020.11.008

46. Cowan DB, Yao R, Thedsanamoorthy JK, Zurakowski D, del Nido PJ, McCully JD. Transit and integration of extracellular mitochondria in human heart cells. Sci Rep. 2017;7(1):17450. doi:10.1038/s41598-017-17813-0

47. Borcherding N, Jia W, Giwa R, et al. Dietary lipids inhibit mitochondria transfer to macrophages to divert adipocyte-derived mitochondria into the blood. Cell Metab. 2022;34(10):1499–1513.e8. doi:10.1016/j.cmet.2022.08.010

48. McCully JD, Cowan DB, Pacak CA, Toumpoulis IK, Dayalan H, Levitsky S. Injection of isolated mitochondria during early reperfusion for cardioprotection. Am J Physiol Heart Circ Physiol. 2009;296(1):H94–H105. doi:10.1152/ajpheart.00567.2008

49. Emani SM, Piekarski BL, Harrild D, Del Nido PJ, McCully JD. Autologous mitochondrial transplantation for dysfunction after ischemia-reperfusion injury. J Thorac Cardiovasc Surg. 2017;154(1):286–289. doi:10.1016/j.jtcvs.2017.02.018

50. Norat P, Sokolowski JD, Gorick CM, et al. Intraarterial Transplantation of Mitochondria After Ischemic Stroke Reduces Cerebral Infarction. Stroke Vasc Interv Neurol. 2023;3(3):e000644. doi:10.1161/svin.122.000644

51. Li SJ, Zheng QW, Zheng J, et al. Mitochondria transplantation transiently rescues cerebellar neurodegeneration improving mitochondrial function and reducing mitophagy in mice. Nat Commun. 2025;16:2839. doi:10.1038/s41467-025-58189-4

52. Lin RZ, Im GB, Luo AC, et al. Mitochondrial transfer mediates endothelial cell engraftment through mitophagy. Nature. 2024;629(8012):660–668. doi:10.1038/s41586-024-07340-0

53. Fisher CR, Ferrington DA. Perspective on AMD Pathobiology: A Bioenergetic Crisis in the RPE. Invest Ophthalmol Vis Sci. 2018;59(4):AMD41–AMD47. doi:10.1167/iovs.18-24289

54. Karunadharma PP, Nordgaard CL, Olsen TW, Ferrington DA. Mitochondrial DNA damage as a potential mechanism for age-related macular degeneration. Invest Ophthalmol Vis Sci. 2010;51(11):5470–5479. doi:10.1167/iovs.10-5429

55. Brown EE, DeWeerd AJ, Ildefonso CJ, Lewin AS, Ash JD. Mitochondrial oxidative stress in the retinal pigment epithelium (RPE) led to metabolic dysfunction in both the RPE and retinal photoreceptors. Redox Biol. 2019;24:101201. doi:10.1016/j.redox.2019.101201

56. Kanow MA, Giarmarco MM, Jankowski CS, et al. Biochemical adaptations of the retina and retinal pigment epithelium support a metabolic ecosystem in the vertebrate eye. Elife. 2017;6:e28899. doi:10.7554/eLife.28899

57. Thiele S, Wu Z, Isselmann B, Pfau M, Guymer RH, Luu CD. Natural History of the Relative Ellipsoid Zone Reflectivity in Age-Related Macular Degeneration. Ophthalmol Retina. 2022;6(12):1165–1172. doi:10.1016/j.oret.2022.06.001

58. Kowluru RA, Chan PS. Oxidative stress and diabetic retinopathy. Exp Diabetes Res. 2007;2007:43603. doi:10.1155/2007/43603

59. Biallosterski C, van Velthoven MEJ, Michels RPJ, Schlingemann RO, DeVries JH, Verbraak FD. Decreased optical coherence tomography-measured pericentral retinal thickness in patients with diabetes mellitus type 1 with minimal diabetic retinopathy. Br J Ophthalmol. 2007;91(9):1135–1138. doi:10.1136/bjo.2006.111534

60. van Dijk HW, Kok PHB, Garvin M, et al. Selective loss of inner retinal layer thickness in type 1 diabetic patients with minimal diabetic retinopathy. Invest Ophthalmol Vis Sci. 2009;50(7):3404–3409. doi:10.1167/iovs.08-3143

61. Yang JH, Kwak HW, Kim TG, Han J, Moon SW, Yu SY. Retinal Neurodegeneration in Type II Diabetic Otsuka Long-Evans Tokushima Fatty Rats. Invest Ophthalmol Vis Sci. 2013;54(6):3844–3851. doi:10.1167/iovs.12-11309

62. Barber AJ, Lieth E, Khin SA, Antonetti DA, Buchanan AG, Gardner TW. Neural apoptosis in the retina during experimental and human diabetes. Early onset and effect of insulin. J Clin Invest. 1998;102(4):783–791. doi:10.1172/JCI2425

63. Sokol S, Moskowitz A, Skarf B, Evans R, Molitch M, Senior B. Contrast sensitivity in diabetics with and without background retinopathy. Arch Ophthalmol. 1985;103(1):51–54. doi:10.1001/archopht.1985.01050010055018

64. Roy MS, Gunkel RD, Podgor MJ. Color vision defects in early diabetic retinopathy. Arch Ophthalmol. 1986;104(2):225–228. doi:10.1001/archopht.1986.01050140079024

65. Cho NC, Poulsen GL, Ver Hoeve JN, Nork TM. Selective loss of S-cones in diabetic retinopathy. Arch Ophthalmol. 2000;118(10):1393–1400. doi:10.1001/archopht.118.10.1393

66. Du Y, Veenstra A, Palczewski K, Kern TS. Photoreceptor cells are major contributors to diabetes-induced oxidative stress and local inflammation in the retina. Proc Natl Acad Sci U S A. 2013;110(41):16586–16591. doi:10.1073/pnas.1314575110

67. Arden GB. The absence of diabetic retinopathy in patients with retinitis pigmentosa: implications for pathophysiology and possible treatment. Br J Ophthalmol. 2001;85(3):366–370. doi:10.1136/bjo.85.3.366

68. de Gooyer TE, Stevenson KA, Humphries P, Simpson DAC, Gardiner TA, Stitt AW. Retinopathy is reduced during experimental diabetes in a mouse model of outer retinal degeneration. Invest Ophthalmol Vis Sci. 2006;47(12):5561–5568. doi:10.1167/iovs.06-0647

69. Kanwar M, Chan PS, Kern TS, Kowluru RA. Oxidative damage in the retinal mitochondria of diabetic mice: possible protection by superoxide dismutase. Invest Ophthalmol Vis Sci. 2007;48(8):3805–3811. doi:10.1167/iovs.06-1280

70. Masser DR, Otalora L, Clark NW, Kinter MT, Elliott MH, Freeman WM. Functional changes in the neural retina occur in the absence of mitochondrial dysfunction in a rodent model of diabetic retinopathy. J Neurochem. 2017;143(5):595–608. doi:10.1111/jnc.14216

71. Madsen-Bouterse SA, Mohammad G, Kanwar M, Kowluru RA. Role of mitochondrial DNA damage in the development of diabetic retinopathy, and the metabolic memory phenomenon associated with its progression. Antioxid Redox Signal. 2010;13(6):797–805. doi:10.1089/ars.2009.2932

72. Miller DJ, Cascio MA, Rosca MG. Diabetic Retinopathy: The Role of Mitochondria in the Neural Retina and Microvascular Disease. Antioxidants (Basel). 2020;9(10):905. doi:10.3390/antiox9100905

73. Kowluru RA. Diabetic retinopathy, metabolic memory and epigenetic modifications. Vision Res. 2017;139:30–38. doi:10.1016/j.visres.2017.02.011

74. Mishra M, Kowluru RA. Epigenetic Modification of Mitochondrial DNA in the Development of Diabetic Retinopathy. Invest Ophthalmol Vis Sci. 2015;56(9):5133–5142. doi:10.1167/iovs.15-16937

75. Takihara Y, Inatani M, Eto K, et al. In vivo imaging of axonal transport of mitochondria in the diseased and aged mammalian CNS. Proc Natl Acad Sci USA. 2015;112(33):10515–10520. doi:10.1073/pnas.1509879112

76. Kimball EC, Pease ME, Steinhart MR, et al. A mouse ocular explant model that enables the study of living optic nerve head events after acute and chronic intraocular pressure elevation: Focusing on retinal ganglion cell axons and mitochondria. Experimental Eye Research. 2017;160:106–115. doi:10.1016/j.exer.2017.04.003

77. Andrews R, Griffiths P, Johnson M, Turnbull D. Histochemical localisation of mitochondrial enzyme activity in human optic nerve and retina. Br J Ophthalmol. 1999;83(2):231–235.

78. Quigley HA, Addicks EM. Chronic experimental glaucoma in primates. II. Effect of extended intraocular pressure elevation on optic nerve head and axonal transport. Invest Ophthalmol Vis Sci. 1980;19(2):137–152.

79. Buckingham BP, Inman DM, Lambert W, et al. Progressive ganglion cell degeneration precedes neuronal loss in a mouse model of glaucoma. J Neurosci. 2008;28(11):2735–2744. doi:10.1523/JNEUROSCI.4443-07.2008

80. Aharoni-Simon M, Ben-Yaakov K, Sharvit-Bader M, et al. Oxidative stress facilitates exogenous mitochondria internalization and survival in retinal ganglion precursor-like cells. Sci Rep. 2022;12(1):5122. doi:10.1038/s41598-022-08747-3

81. Noh SE, Lee SJ, Lee TG, Park KS, Kim JH. Inhibition of cellular senescence hallmarks by mitochondrial transplantation in senescence-induced ARPE-19 cells. Neurobiol Aging. 2023;121:157–165. doi:10.1016/j.neurobiolaging.2022.11.003

82. Noh SE, Lee SJ, Cho CS, Jo DH, Park KS, Kim JH. Mitochondrial transplantation attenuates oligomeric amyloid-beta-induced mitochondrial dysfunction and tight junction protein destruction in retinal pigment epithelium. Free Radic Biol Med. 2024;212:10–21. doi:10.1016/j.freeradbiomed.2023.12.012

83. Ashok A, Chen DF. Mitochondria Transplantation Preserves Retinal Ganglion Cells and Promotes CNS Axonal. Social Science Research Network. Preprint posted online December 19, 2025:5929018. doi:10.2139/ssrn.5929018

84. Nascimento-Dos-Santos G, de-Souza-Ferreira E, Lani R, et al. Neuroprotection from optic nerve injury and modulation of oxidative metabolism by transplantation of active mitochondria to the retina. Biochim Biophys Acta Mol Basis Dis. 2020;1866(5):165686. doi:10.1016/j.bbadis.2020.165686

85. Wu SF, Lin CY, Tsai RK, et al. Mitochondrial Transplantation Moderately Ameliorates Retinal Degeneration in Royal College of Surgeons Rats. Biomedicines. 2022;10(11):2883. doi:10.3390/biomedicines10112883

86. Wang Y, Liu N, Hu L, et al. Nanoengineered mitochondria enable ocular mitochondrial disease therapy via the replacement of dysfunctional mitochondria. Acta Pharm Sin B. 2024;14(12):5435–5450. doi:10.1016/j.apsb.2024.08.007

87. Garweg JG, Keiper J, Pfister IB, Schild C. Functional Outcomes of Brolucizumab-Induced Intraocular Inflammation Involving the Posterior Segment-A Meta-Analysis and Systematic Review. J Clin Med. 2023;12(14):4671. doi:10.3390/jcm12144671

88. Wykoff CC, Matsumoto H, Barakat MR, et al. RETINAL VASCULITIS OR VASCULAR OCCLUSION AFTER BROLUCIZUMAB FOR NEOVASCULAR AGE-RELATED MACULAR DEGENERATION. Retina. 2023;43(7):1051–1063. doi:10.1097/IAE.0000000000003769

89. Nahar A, Eliott D, Avery RL, et al. Retinal Vasculopathy and Choroiditis after Pegcetacoplan Injection: Clinicopathologic Support for a Drug Hypersensitivity Reaction. Ophthalmol Retina. 2025;9(4):352–366. doi:10.1016/j.oret.2024.10.005

90. Weiß E, Kretschmer D. Formyl-Peptide Receptors in Infection, Inflammation, and Cancer. Trends Immunol. 2018;39(10):815–829. doi:10.1016/j.it.2018.08.005

91. Wu G, Zhu Q, Zeng J, et al. Extracellular mitochondrial DNA promote NLRP3 inflammasome activation and induce acute lung injury through TLR9 and NF-κB. J Thorac Dis. 2019;11(11):4816–4828. doi:10.21037/jtd.2019.10.26

92. Emani SM, Piekarski BL, Harrild D, Del Nido PJ, McCully JD. Autologous mitochondrial transplantation for dysfunction after ischemia-reperfusion injury. J Thorac Cardiovasc Surg. 2017;154(1):286–289. doi:10.1016/j.jtcvs.2017.02.018

93. Wu KY, Gao A, Giunta M, Tran SD. What’s New in Ocular Drug Delivery: Advances in Suprachoroidal Injection since 2023. Pharmaceuticals (Basel). 2024;17(8):1007. doi:10.3390/ph17081007

94. Patel MJ, Sheth S, Mar J, Gregori NZ, Sengillo JD. Surgical Approaches to Retinal Gene Therapy: 2025 Update. Bioengineering (Basel). 2025;12(10):1122. doi:10.3390/bioengineering12101122

95. Mahajan VB, Barak Y, Almeida DRP. Near-Real Surgical Specimens (NRSS): A Novel Platform for Standardized Assessment of Suprachoroidal Drug Delivery. Transl Vis Sci Technol. 2025;14(9):6. doi:10.1167/tvst.14.9.6

96. Barak Y, Tamir KM, Adakhovska A, Mangelus M, Hoggeg AB. First-In-Human Results of a Novel Suprachoroidal Delivery Injector. Invest Ophthalmol Vis Sci. 2025;66(8):1774–1774.

97. Rotenstreich Y, Sher I, Lawrence M, Mangelus M, Ingerman A, Barak Y. A Novel Device for Suprachoroidal Drug Delivery to Retina: Evaluation in Nonhuman Primates. Transl Vis Sci Technol. 2023;12(6):3. doi:10.1167/tvst.12.6.3

98. Melki I, Allaeys I, Tessandier N, et al. Platelets release mitochondrial antigens in systemic lupus erythematosus. Sci Transl Med. 2021;13(581):eaav5928. doi:10.1126/scitranslmed.aav5928

99. Becker Y, Loignon RC, Julien AS, et al. Anti-mitochondrial autoantibodies in systemic lupus erythematosus and their association with disease manifestations. Sci Rep. 2019;9(1):4530. doi:10.1038/s41598-019-40900-3

100. Labitzke EM, Diani-Moore S, Rifkind AB. Mitochondrial P450-dependent arachidonic acid metabolism by TCDD-induced hepatic CYP1A5; conversion of EETs to DHETs by mitochondrial soluble epoxide hydrolase. Archives of biochemistry and biophysics. 2007;468(1):70. doi:10.1016/j.abb.2007.08.012

101. Wert KJ, Skeie JM, Davis RJ, Tsang SH, Mahajan VB. Subretinal Injection of Gene Therapy Vectors and Stem Cells in the Perinatal Mouse Eye. J Vis Exp. 2012;(69):4286. doi:10.3791/4286

