## Supplementary material for "Mitochondrial Transplantation in the Eye: A Review and Evaluation of Surgical Approaches": Table 1-3

**Table 1. In vivo ocular mitochondrial transplantation studies**

| **Study** | **Model** | **Donor mitochondria** | **Delivery** | **Evidence/ Key readouts** | **Main findings** |
| --- | --- | --- | --- | --- | --- |
| Nascimento-dos-Santos et al., 2020 | Rat ONC | Rat liver mitochondria (MitoTracker) | IVT, single | Confocal imaging, Seahorse assay, ERG, CTB axon tracing | Transplanted mitochondria improved retinal respiration, increased TUJ1+ RGC survival, and promoted axon regeneration beyond the crush site; effects were lost with lysed/depolarized mitochondria. |
| Wu et al., 2022 | RCS rat retinal degeneration | Rat liver mitochondria (unlabeled) | IVT, weekly x5 | OCT, VEP, H&E/ONL thickness, TUNEL | Repeated mitochondrial delivery produced partial structural and functional rescue, including reduced retinal thinning, improved ONL preservation, and partial VEP maintenance. |
| Noh et al., 2024 | Mouse subretinal FITC‑oAβ | Human bone marrow-mononuclear cell line (unlabeled) | IVT, single | Confocal imaging with human-specific anti-mitochondrial staining, retinal sections, RPE flatmounts, ZO-1/phalloidin, FITC-oAβ clearance | Donor mitochondria were detected across retinal layers including RPE, with reduced oAβ burden and improved RPE tight junction architecture, supporting bioenergetic rescue and barrier protection. |
| Wang et al., 2024 | Rotenone LHON-like mouse) | Heart mitochondria+ PARKIN mRNA‑nanoparticles (MitoTracker) | IVT, single | Confocal retinal localization, PARKIN/mitophagy markers, complex I activity, ATP, H&E, cytokines, mtDNA safety assays, optomotor testing | Combined mitochondrial supplementation and PARKIN-driven mitophagy increased ATP and complex I activity, improved retinal structure, and enhanced visual behavior. |
| Ashok et al., 2025 (preprint) | Mouse ONC | P3 mouse liver mitochondria (MitoTracker) | IVT, single | Confocal imaging, RBPMS RGC counts, ERG, anterograde axon tracing, TEM | Donor mitochondria localized to the RGC layer and were associated with increased RGC survival, improved ERG, enhanced axon regeneration, and improved mitochondrial ultrastructure in regenerating axons. |

*Abbreviations: ATP, adenosine triphosphate; CTB, cholera toxin subunit B; ERG, electroretinography; FITC, fluorescein isothiocyanate; IVT, intravitreal injection; LHON, Leber hereditary optic neuropathy; mtDNA, mitochondrial DNA; oAβ, oligomeric amyloid-beta; ONC, optic nerve crush; ONL, outer nuclear layer; OCT, optical coherence tomography; PARKIN, parkin RBR E3 ubiquitin protein ligase; RBPMS, RNA-binding protein with multiple splicing; RCS, Royal College of Surgeons; RGC, retinal ganglion cell; TEM, transmission electron microscopy; VEP, visual evoked potential; ZO-1, zonula occludens-1.*

**Table 2. In vitro ocular mitochondrial transplantation studies**

| **Study** | **Model** | **Donor mitochondria** | **Evidence/ Key readouts** | **Evidence/ Key readouts** |
| --- | --- | --- | --- | --- |
| Aharoni-Simon et al., 2022 | 661W/oxidative stress (H2O2) | Mouse liver mitochondria (MitoTracker) | Flow cytometry, confocal imaging, strain-specific mtDNA qPCR | Exogenous mitochondria showed dose-dependent uptake; oxidative stress enhanced uptake and prolonged donor mitochondrial retention up to 72 h. |
| Noh et al., 2023 | ARPE-19 senescence (replicative/oxidative) | UC-MSC mitochondria (MitoTracker) | Confocal imaging, SA-β-gal, p16/p21, NF-κB activity, inflammatory cytokines, mitochondrial function/dynamics assays | Mitochondrial transfer reduced senescence and inflammatory markers and improved mitochondrial function and quality-control signaling. |
| Noh et al., 2024 | ARPE-19 amyloid-beta stress | MSC mitochondria (MitoTracker) | Confocal imaging, Δψm, ATP, ROS, ZO-1/tight junction proteins, barrier integrity, oAβ clearance assays | Donor mitochondria alleviated bioenergetic dysfunction, preserved tight junction/barrier integrity, and improved oAβ clearance. |
| Wang et al., 2024 | HeLA rotenone-induced complex I injury | Heart mitochondria + PARKIN mRNA nanoparticles (MitoTracker) | Confocal imaging, complex I activity/protein, ATP, ROS, mitophagy markers | Nanoengineered mitochondria improved uptake, enhanced PARKIN-mediated mitophagy, and restored complex I function and ATP production. |
| Ashok et al., 2025 (preprint) | SH-SY5Y glutamate injury; PC12 trophic deprivation; primary RGC culture | P3 mouse liver (MitoTracker) | Confocal imaging, neurite length/number, JC-1, Δψm, ROS, ATP, βIII-tubulin, GAP43 | Mitochondrial supplementation improved cell survival and neurite outgrowth, restored mitochondrial membrane potential and ATP, and reduced ROS in stressed neuronal cells. |

*Abbreviations: Aβ, amyloid-beta; ARPE-19, human retinal pigment epithelial cell line-19; ATP, adenosine triphosphate; Δψm, mitochondrial membrane potential; H2O2, hydrogen peroxide; MSC, mesenchymal stromal/stem cell; mtDNA, mitochondrial DNA; qPCR, quantitative polymerase chain reaction; RGC, retinal ganglion cell; ROS, reactive oxygen species; SH-SY5Y, human neuroblastoma cell line; ZO-1, zonula occludens-1.*

**Table 3. Clinical and Biological Considerations of Subretinal, Intravitreal, and Suprachoroidal Delivery**

| **Feature** | **Subretinal Injection** | **Intravitreal Injection** | **Suprachoroidal Injection** |
| --- | --- | --- | --- |
| **Primary Localization** | RPE, ONL | Inner retina (GCL, IPL, INL), Optic nerve fibers | Suprachoroidal Space (Broad distribution in posterior segment, outer-retina biased distribution) |
| **Surgical Complexity** | High (Requires operating room, complex maneuvers) | Low (Routine clinical procedure) | Moderate/Low (Minimally invasive, potentially office-based) |
| **Injection Volume** | Moderate (~0.05–0.15 mL), confined to a localized bleb | Low (0.05–0.1 mL), due to IOP limits | Moderate to High (~0.1 mL), can accommodate larger volumes for broad distribution |
| **Key Biological Barrier** | ELM | Vitreous matrix, ILM | Bruch’s Membrane |
| **Proposed Mechanism** | Phagocytosis by RPE | Endocytosis by inner retinal cells; possible axonal transport when RGCs are targeted | Trans-tissue diffusion with secondary cellular uptake |
| **Clinical Translatability** | Limited by risk of retinal detachment and foveal damage; difficult to repeat | Highly scalable, but limited by low concentration reaching the outer retina | Promising for repeated dosing; avoids retinal detachment risks |

*Abbreviations: ELM, external limiting membrane; GCL, ganglion cell layer; ILM, internal limiting membrane; INL, inner nuclear layer; IOP, intraocular pressure; IPL, inner plexiform layer; ONL, outer nuclear layer; RPE, retinal pigment epithelium.*
